# Genomic sequencing and neutralizing serological profiles during acute dengue infection: A 2017 cohort study in Nepal

**DOI:** 10.1101/2024.06.03.597174

**Authors:** Sabita Prajapati, Annie Elong Ngono, Melanie Mc Cauley, Julia Timis, Srijan Shrestha, Anup Bastola, Shrawan Kumar Mandal, Sanjay Ray Yadav, Rajindra Napit, Meng Ling Moi, Montarop Yamabhai, October M Sessions, Sujan Shresta, Krishna Das Manandhar

**Author notes:** Corresponding authors (KDM), (OMS) and (SS). Emerging Infectious Disease Branch, Walter Reed Army Institute of Research, Silver Spring, Maryland, USA. These authors contributed equally (Sabita Prajapati, Annie Elong Ngono).

## Abstract

Dengue virus (DENV) is a mosquito-borne flavivirus that poses a threat to nearly 50% of the global population. DENV has been endemic in Nepal since 2006; however, little is known about how DENV is evolving or the prevalence of anti-DENV immunity within the Nepalese population. To begin to address these gaps, we performed a serologic and genetic study of 49 patients from across Nepal who presented at central hospitals during the 2017 dengue season with suspected DENV infection. Of the 49 subjects assessed, 21 (43%) were positive for DENV NS1 antigen; of these; 5 were also anti-DENV IgM^+^ IgG^+^; 7 were DENV IgM^+^ IgG^−^, 2 were IgM^−^ IgG^+^, and 7 were IgM^-^ IgG^−^ by specific ELISAs. Seven of the 21 NS1+ sera were RNA+ by RT-PCR (six DENV2, one DENV3), suggesting that DENV2 was the dominant serotype in our cohort. Whole-genome sequencing of two DENV2 isolates showed similarity with strains circulating in Singapore in 2016, and the envelope genes were also similar to strains circulating in India in 2017. DENV-neutralizing antibodies (nAbs) were present in 31 of 47 sera tested (66%); among these, 20, 24, 26, and 12 sera contained nAbs against DENV1, 2, 3, and 4 serotypes, respectively. Serology analysis suggested that 12 (26%) and 19 (40%) of the 49 subjects were experiencing primary and secondary DENV infections, respectively. Collectively, our results provide evidence for current and/or past exposure to multiple DENV serotypes in our cohort, and the RNA analyses further indicate that DENV2 was the likely dominant serotype circulating in Nepal in 2017. These data suggest that expanded local surveillance of circulating DENV genotypes and population immunity will be important to effectively manage and mitigate future dengue outbreaks in Nepal.

## Introduction

Dengue virus (DENV) is a positive-sense single-stranded RNA virus of the *Flavivirus* genus, which also includes Japanese encephalitis, Zika, yellow fever, and West Nile viruses (1, 2)—all transmitted, in the vast majority of cases, by infected *Aedes* spp. mosquitoes (3, 4). The four antigenically distinct serotypes of DENV (DENV1–4) are responsible for roughly 400 million reported DENV infections per year (3, 4), which may be asymptomatic or have symptoms ranging from self-limiting dengue fever to severe disease characterized by hemorrhagic fever, shock and, in some cases, death (1).

Infection with DENV confers long-term protection against homologous serotypes but only limited cross-protection against heterologous serotypes; indeed, in some cases, secondary infection with a different serotype can elicit severe dengue in an antibody (Ab)-dependent manner (5-7). At present, there are no DENV vaccines that provide durable protection against all four DENV serotypes and can be administered to people of all ages who are DENV-naive or DENV-immune. The first approved dengue vaccine (DengVaxia) can only be administered to individuals with prior natural DENV infection (8), because it increases the risk of severe dengue in DENV-naive individuals who are then exposed naturally after vaccination. A new vaccine, QDENGA (Takeda) has been recently approved by Europe, Indonesia, Thailand, Argentina, and Brazil for DENV-naive and -immune individuals, but it provides robust long-term protection against only selected DENV serotypes (9-11). Finally, a phase 3 trial of the tetravalent vaccine Butantan-DV (Instituto Butantan/NIAID) demonstrated good protection against DENV1, but was only moderately effective against DENV2 (12). These data highlight the challenges in developing a DENV vaccine that induces protective immunity against all four serotypes regardless of prior natural exposure. Moreover, the potential dangers of vaccination mean that data on currently circulating DENV serotypes and population immune status must be available to enable governments to make informed decisions in selecting the optimal vaccine for distribution in any given year (13).

In Nepal, the first DENV infection was detected in 2004 in a Japanese traveler, from whom DENV2 was isolated (14); by 2006, all 4 DENV serotypes had been documented in Nepal (15). Since then, sporadic cases have been reported during the annual dengue season (September to December), interrupted by major outbreaks roughly every 3 years beginning in 2010 (16). Notably, each successive outbreak has been accompanied by an increase in case numbers, geographic spread, morbidity, and mortality (17-19). In 2017, a total of 2111 dengue cases were reported from 28 of Nepal’s 77 districts whereas in 2019, 17,992 cases, including 6 deaths, were reported in 68 of the 77 districts (16, 20, 21), representing a 140-fold increase in incidence in just 2 years (16). This trend continued in the 2022 outbreak, with 54,784 reported cases and 88 deaths involving all districts (17). Based on these data, Nepal is on a trajectory to experience another DENV outbreak in 2025.

To date, very little data have been collected on DENV virology and immunology in the Nepalese population. In part, this is because Nepal is a low-income country with a limited scientific infrastructure. For example, the full genomes of only few circulating DENV isolates have been sequenced since the first reported infection in Nepal in 2006 (22). The present study was conducted as part of a long-term process of enabling local scientists/clinicians to effectively surveil the circulation of, and population exposure to, DENV serotypes in Nepal. To this end, we describe here serological and genomic analyses of sera from 49 residents of districts across Nepal who presented with suspected dengue at hospitals in Kathmandu and Chitwan, two geographically and climatically distinct districts. All sera were analyzed for DENV-NS1 antigen, DENV-specific anti-IgM/IgG, and neutralizing antibody (nAb) response against all four DENV serotypes. A subset of NS1+ sera were further serotyped by RT-PCR and subjected to whole-genome sequencing. The knowledge gained from this study has already facilitated the expansion of DENV genomic surveillance studies by local scientists/clinicians with the goal of mitigating the dengue outbreak predicted to occur in 2025.

## Materials and methods

### Sample collection and ethical approval

The study cohort consisted of 49 febrile patients who presented with suspected dengue at Sukraraj Tropical and Infectious Disease Hospital in Kathmandu and at Chitwan Medical College and Teaching Hospital in Chitwan during the 2017 dengue outbreak (September 2017 through January 2018). A diagnosis of suspected dengue was made by attending physicians. Venous blood samples (5 mL) were collected into EDTA tubes, and demographic and clinicopathological information was recorded. Ethical approval was obtained from the Nepal Health Research Council (Reg. no. 378/2016). Written informed consent was obtained from either the adult patient or a parent/guardian for patients under 18 years of age. All steps were performed by the Nepal-based authors and teams, with advice from the authors based in the US, Japan, and Thailand.

### Enzyme-linked immunosorbent assays

Plasma was isolated from the blood samples and stored at -80°C until analyzed. Levels of DENV NS1 antigen, anti-DENV IgM, and anti-DENV IgG were quantified using commercial ELISA kits (InBios: DNS1-R, DDMS-1 and DDGS-R respectively). Assay procedures, calculation of immune status ratios, and classification as seronegative or seropositive were all performed according to the manufacturer’s instructions. ELISAs were performed at the Infectious and Viral Disease Research Laboratory, Central Department of Biotechnology, Tribhuvan University in Nepal in the year 2018.

### DENV serotyping by RT-PCR

Viral RNA was isolated from plasma samples using a QIAamp Viral RNA Mini kit (Qiagen, 52906). DENV1–4 serotypes were identified by multiplex RT-PCR as described previously (23-26), using US Centers for Disease Control and Prevention real-time RT-PCR assay kits (KK0128) at the La Jolla Institute for Immunology in 2018.

### Whole genome DENV sequencing

All procedures were conducted according to the kit manufacturer’s recommendations. Illumina libraries were constructed from total RNA using the NEB Next Ultra Directional RNA Library Prep Kit (New England Biolabs, E7760) per manufacturer’s instructions. Libraries of 400–600 nucleotides were obtained using Mag-Bind RxnPure Plus beads (Omega Bio-Tek, M1386-01), purified with the MinElute PCR Purification Kit (Qiagen, 28004), and quantified using a Bioanalyzer High-Sensitivity DNA Assay (Agilent Technologies, 5067-4626). Targeted DENV genome enrichment was achieved using custom-designed biotinylated 120-mer xGen Lockdown baits (Integrated DNA Technologies) with complementarity to DENV1–4 serotypes, as previously described (27), all per manufacturer’s instructions. Genome assembly was performed using the VIPR4 pipeline (https://github.com/nf-core/vipr/). Whole genome sequencing was performed and analyzed at the National University of Singapore in 2019.

### Phylogenetic analysis

Multiple sequence alignment of DENV2 full genome sequences from the present study (n=2) and the National Center for Biotechnology Information (NCBI) (n=1,777) was carried out using a fast Fourier transformation method in MAFFT v7.490. An approximately maximum likelihood phylogenetic tree was generated using a generalized time-reversible model of nucleotide evolution in FastTree v2.1.11, uses SH-like local supports with 1,000 resamples to estimate and validate the reliability of each split in the tree. The branch containing the 2 sequences from the 2017 Nepal outbreak and 29 complete genome sequences from the NCBI were selected, and a more robust maximum likelihood phylogenetic tree created using RAXML v8.2.11 and the GTR GAMMA model with 1,000 bootstrap replications. Trees were visualized using FigTree v1.4.4.

### Flow cytometry-based DENV1–4 neutralization assay

Neutralization assays were performed as previously described (28), using a U937 DC-SIGN cell-based flow cytometry assay. Briefly, plasma samples were serially diluted 5-fold starting at 1:40, and incubated with pre-titrated DENV1-4 serotypes (WHO reference standards) for 1 h at 37°C. U937 DC-SIGN cells were then incubated with the plasma/virus mixtures for 2 h at 37°C, washed, and incubated again for 16 h at 37°C. As controls, cells were incubated with virus in the absence of serum to obtain the baseline infection rate. The cells were then surface stained with anti-CD209-PE Ab (DC-SIGN) and intracellularly stained with anti-FITC-labeled 4G2 Ab (pan-flaviviral envelope [E] protein), and analyzed using a FACScanto LSR cytometer and FlowJo V9 software (BD Biosciences).

Serum was considered positive for neutralizing activity if DENV infection was reduced by ≥90% at a serum dilution of >1:40 (cutoff = NT_90_ >40). NT_90_ values were used to classify primary infection (first infection with a single serotype) or secondary infection (prior infections with other serotypes), as previously described (29). In brief, primary infection was defined as (i) an NT_90_ ≥40 against a single DENV serotype and <40 against all other serotypes, or (ii) a response against a single DENV serotype, with an NT_90_ 4-fold higher than against the other serotypes. Secondary infection was defined as (i) an NT_90_ ≥40 against at least two DENV serotypes, and/or (ii) a response with an NT_90_ ≤4-fold higher than the next highest response (29). This assay and analysis were performed at the La Jolla Institute for Immunology in 2018 and 2019.

## Results

### Demography and disease association with platelet count

The study cohort consisted of 49 patients who presented with suspected dengue at two major hospitals in Kathmandu and Chitwan, Nepal during the 2017 dengue season (September 2017 to January 2018) (30). The majority of the subjects were male (71%, 35/49) and were aged 20 to 59 years (63%, 31/49), with a median age of 34 years (range 13–81) (**Table 1**). Although the 49 subjects presented to the two major hospitals in Kathmandu and Chitwan, they originated from 6 of Nepal’s 7 provinces (**Fig 1**).

**Table 1.**
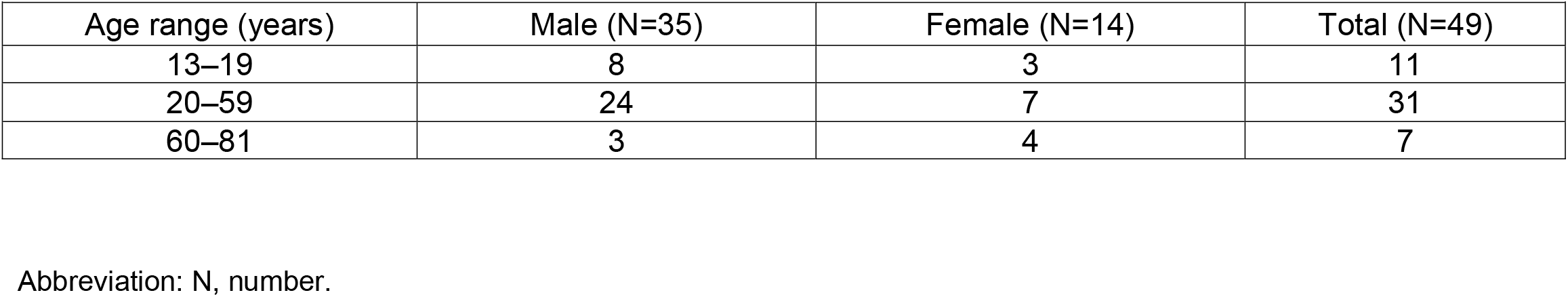
Characteristics of study participants.

**Fig 1.**
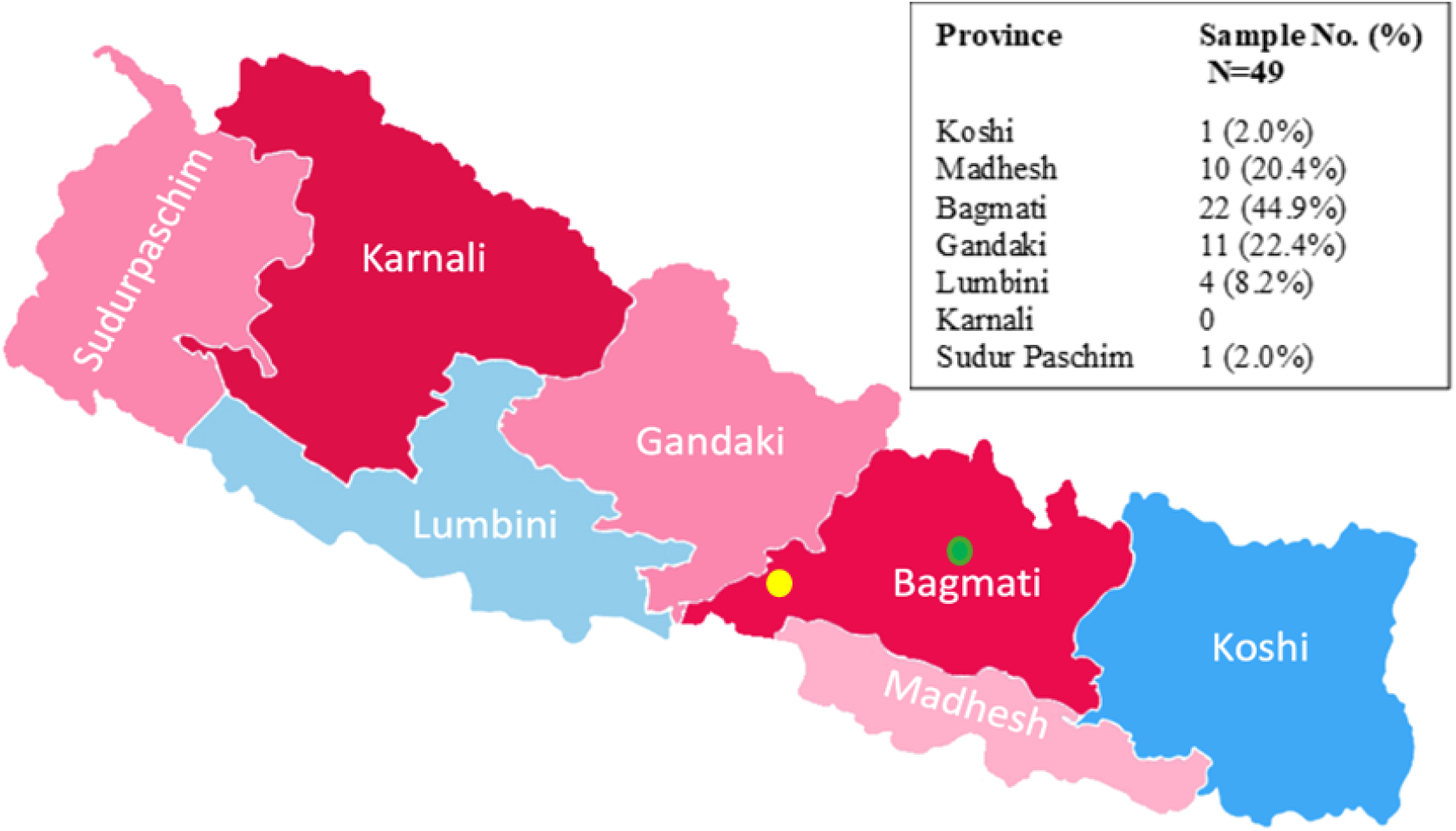
Geographic distribution of the home residences of the study subjects. Patients originated from across Nepal and were seen at hospitals in Kathmandu (green dot) and Chitwan (yellow dot). Abbreviation: No, number.

All subjects were symptomatic at the time of sample collection, although the time between symptom onset and blood collection was not recorded. Thrombocytopenia is a potential indicator of dengue disease severity (31), and platelet counts of 41–100 x 10^3^/μL, 21–40 x 10^3^/μL, and ≤20 × 10^3^/μL are considered to reflect low, moderate, and high risk, respectively, of bleeding associated with severe dengue (normal platelet counts: 100– 450 x 10^3^/μL) (32). Most samples (31/48) had normal platelet counts, with 13, 2, and 2 samples having counts within the low-, moderate-, and high-risk categories, respectively (**Table 2**). These results are consistent with the majority of our study cohort having mild dengue disease.

**Table 2.**
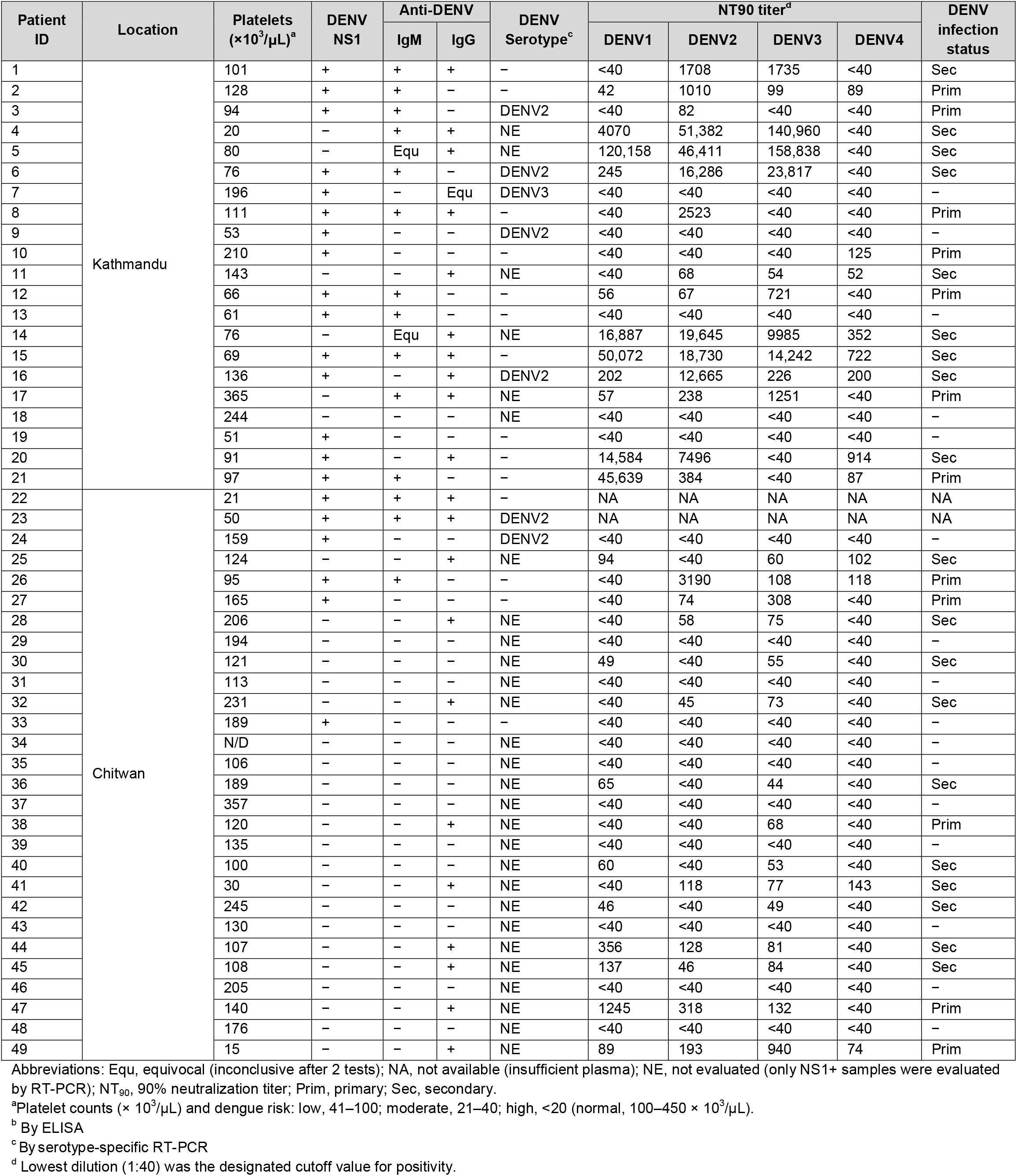
Platelet counts and DENV serology, serotyping, neutralization titers, and infection status.

| Patient ID | Location | Platelets (×10<sup>3</sup>/μL)<sup>a</sup> | DENV NS1 | Anti-DENV |  | DENV Serotype<sup>c</sup> | NT90 titer<sup>d</sup> |  |  |  | DENV infection status |
| --- | --- | --- | --- | --- | --- | --- | --- | --- | --- | --- | --- |
|  |  |  |  | IgM | IgG |  | DENV1 | DENV2 | DENV3 | DENV4 |  |
| 1 | Kathmandu | 101 | + | + | + | – | <40 | 1708 | 1735 | <40 | Sec |
| 2 |  | 128 | + | + | – | – | 42 | 1010 | 99 | 89 | Prim |
| 3 |  | 94 | + | + | – | DENV2 | <40 | 82 | <40 | <40 | Prim |
| 4 |  | 20 | – | + | + | NE | 4070 | 51,382 | 140,960 | <40 | Sec |
| 5 |  | 80 | – | Equ | + | NE | 120,158 | 46,411 | 158,838 | <40 | Sec |
| 6 |  | 76 | + | + | – | DENV2 | 245 | 16,286 | 23,817 | <40 | Sec |
| 7 |  | 196 | + | – | Equ | DENV3 | <40 | <40 | <40 | <40 | – |
| 8 |  | 111 | + | + | + | – | <40 | 2523 | <40 | <40 | Prim |
| 9 |  | 53 | + | – | – | DENV2 | <40 | <40 | <40 | <40 | – |
| 10 |  | 210 | + | – | – | – | <40 | <40 | <40 | 125 | Prim |
| 11 |  | 143 | – | – | + | NE | <40 | 68 | 54 | 52 | Sec |
| 12 |  | 66 | + | + | – | – | 56 | 67 | 721 | <40 | Prim |
| 13 |  | 61 | + | + | – | – | <40 | <40 | <40 | <40 | – |
| 14 |  | 76 | – | Equ | + | NE | 16,887 | 19,645 | 9985 | 352 | Sec |
| 15 |  | 69 | + | + | + | – | 50,072 | 18,730 | 14,242 | 722 | Sec |
| 16 |  | 136 | + | – | + | DENV2 | 202 | 12,665 | 226 | 200 | Sec |
| 17 |  | 365 | – | + | + | NE | 57 | 238 | 1251 | <40 | Prim |
| 18 |  | 244 | – | – | – | NE | <40 | <40 | <40 | <40 | – |
| 19 |  | 51 | + | – | – | – | <40 | <40 | <40 | <40 | – |
| 20 |  | 91 | + | – | + | – | 14,584 | 7496 | <40 | 914 | Sec |
| 21 |  | 97 | + | + | – | – | 45,639 | 384 | <40 | 87 | Prim |
| 22 | Chitwan | 21 | + | + | + | – | NA | NA | NA | NA | NA |
| 23 |  | 50 | + | + | + | DENV2 | NA | NA | NA | NA | NA |
| 24 |  | 159 | + | – | – | DENV2 | <40 | <40 | <40 | <40 | – |
| 25 |  | 124 | – | – | + | NE | 94 | <40 | 60 | 102 | Sec |
| 26 |  | 95 | + | + | – | – | <40 | 3190 | 108 | 118 | Prim |
| 27 |  | 165 | + | – | – | – | <40 | 74 | 308 | <40 | Prim |
| 28 |  | 206 | – | – | + | NE | <40 | 58 | 75 | <40 | Sec |
| 29 |  | 194 | – | – | – | NE | <40 | <40 | <40 | <40 | – |
| 30 |  | 121 | – | – | – | NE | 49 | <40 | 55 | <40 | Sec |
| 31 |  | 113 | – | – | – | NE | <40 | <40 | <40 | <40 | – |
| 32 |  | 231 | – | – | + | NE | <40 | 45 | 73 | <40 | Sec |
| 33 |  | 189 | + | – | – | – | <40 | <40 | <40 | <40 | – |
| 34 |  | N/D | – | – | – | NE | <40 | <40 | <40 | <40 | – |
| 35 |  | 106 | – | – | – | NE | <40 | <40 | <40 | <40 | – |
| 36 |  | 189 | – | – | – | NE | 65 | <40 | 44 | <40 | Sec |
| 37 |  | 357 | – | – | – | NE | <40 | <40 | <40 | <40 | – |
| 38 |  | 120 | – | – | + | NE | <40 | <40 | 68 | <40 | Prim |
| 39 |  | 135 | – | – | – | NE | <40 | <40 | <40 | <40 | – |
| 40 |  | 100 | – | – | – | NE | 60 | <40 | 53 | <40 | Sec |
| 41 |  | 30 | – | – | + | NE | <40 | 118 | 77 | 143 | Sec |
| 42 |  | 245 | – | – | – | NE | 46 | <40 | 49 | <40 | Sec |
| 43 |  | 130 | – | – | – | NE | <40 | <40 | <40 | <40 | – |
| 44 |  | 107 | – | – | + | NE | 356 | 128 | 81 | <40 | Sec |
| 45 |  | 108 | – | – | + | NE | 137 | 46 | 84 | <40 | Sec |
| 46 |  | 205 | – | – | – | NE | <40 | <40 | <40 | <40 | – |
| 47 |  | 140 | – | – | + | NE | 1245 | 318 | 132 | <40 | Prim |
| 48 |  | 176 | – | – | – | NE | <40 | <40 | <40 | <40 | – |
| 49 |  | 15 | – | – | + | NE | 89 | 193 | 940 | 74 | Prim |

Abbreviations: Equ, equivocal (inconclusive after 2 tests); NA, not available (insufficient plasma); NE, not evaluated (only NS1+ samples were evaluated by RT-PCR); NT<sub>90</sub>, 90% neutralization titer; Prim, primary; Sec, secondary.
<sup>a</sup>Platelet counts (× 10<sup>3</sup>/μL) and dengue risk: low, 41–100; moderate, 21–40; high, <20 (normal, 100–450 × 10<sup>3</sup>/μL).
<sup>b</sup> By ELISA
<sup>c</sup> By serotype-specific RT-PCR
<sup>d</sup> Lowest dilution (1:40) was the designated cutoff value for positivity.

### DENV serostatus and serotypes

NS1 antigen was detected in 43% (21/49) of samples, anti-DENV IgM Ab in 29% (14/49) and anti-DENV IgG Ab in 43% (21/49). Thus, 47% (23/49) of our cohort were considered to have active dengue at the time of sample collection based on the presence of ≥1 of the following 5 criteria: NS1^+^ (14%, n=7), NS1^+^IgM^+^ (14%, n=7), NS1^+^IgG^+^ (4%, n=2), NS1^+^IgM^+^IgG^+^ (10%, n=5) and IgM^+^ IgG^+^ (4%, n=2). Previous dengue cases were observed with only IgG^+^ subjects (24%, n=12) (**Table 2**). Samples from 29% (14) showed no evidence of current or prior infection (NS1^−^, IgM^−^, and IgG^−^).. The 21 NS1^+^ sera were further analyzed for DENV serotype by RT-PCR, 7 (33%) were found to be positive for DENV RNA, with 6 samples serotyped as DENV2 and 1 as DENV3 (**Table 2**). Thus, despite the small number of RT-PCR RNA^+^ positive samples, DENV2 appears to have been the dominant circulating serotype in our 49-subject cohort at the time of the 2017 outbreak.

### Neutralizing Ab response against DENV1–4 serotypes

Of the 49 samples collected, 47 were evaluated in cell-based neutralizing assays (**Fig 2A**). The mean NT_90_ titers for DENV1, DENV2, and DENV3 were similar, whereas the mean NT_90_ titer for DENV4 was much lower overall, and significantly lower when compared with DENV2 (**Fig 2A**). Anti-DENV4 nAbs were undetectable in 35 samples (74%) (**Fig 2A**). Strikingly, 3 of the 7 samples positive for DENV RNA+ by RT-PCR were negative for nAbs against any of the DENV serotypes (**Table 2**). By evaluating anti-DENV nAb activity, it is possible to infer whether DENV infection is primary or secondary for single- and multiple-serotype infections (33). Using this approach, 26% (12/47) and 40% (19/47) of samples were from patients experiencing primary and secondary infections, respectively (**Table 2**). The remaining 34% (16/47) of samples had NT_90_ values <40 for all 4 serotypes; of these, 10 were NS1^−^ IgM^−^ IgG^−^ and an additional 4 were IgM^−^ IgG^−^ (**Table 2**). Thus, our study cohort consisted of patients who experienced both primary and secondary DENV exposure during the 2017 outbreak, with a higher prevalence of secondary infections.

**Fig 2.**
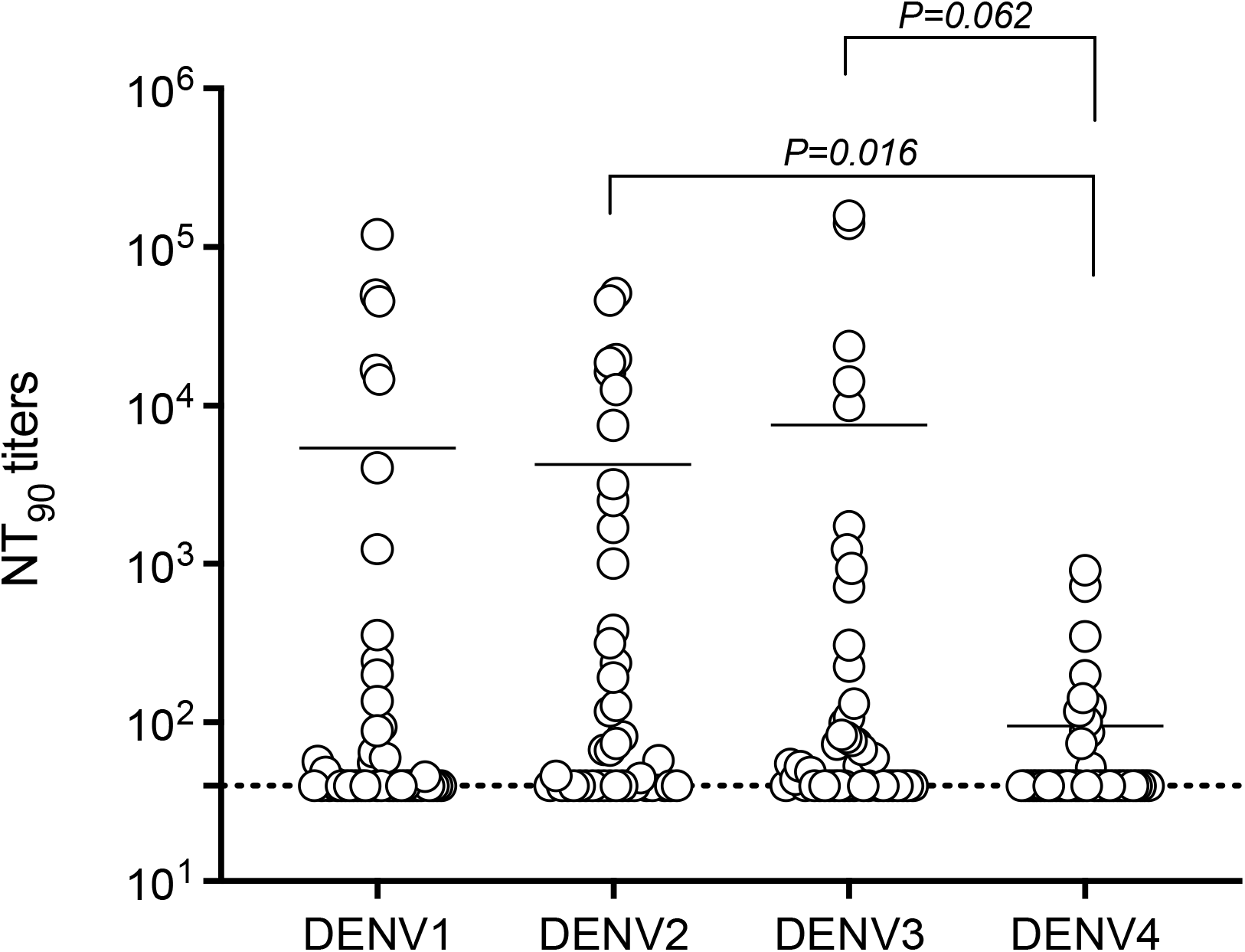
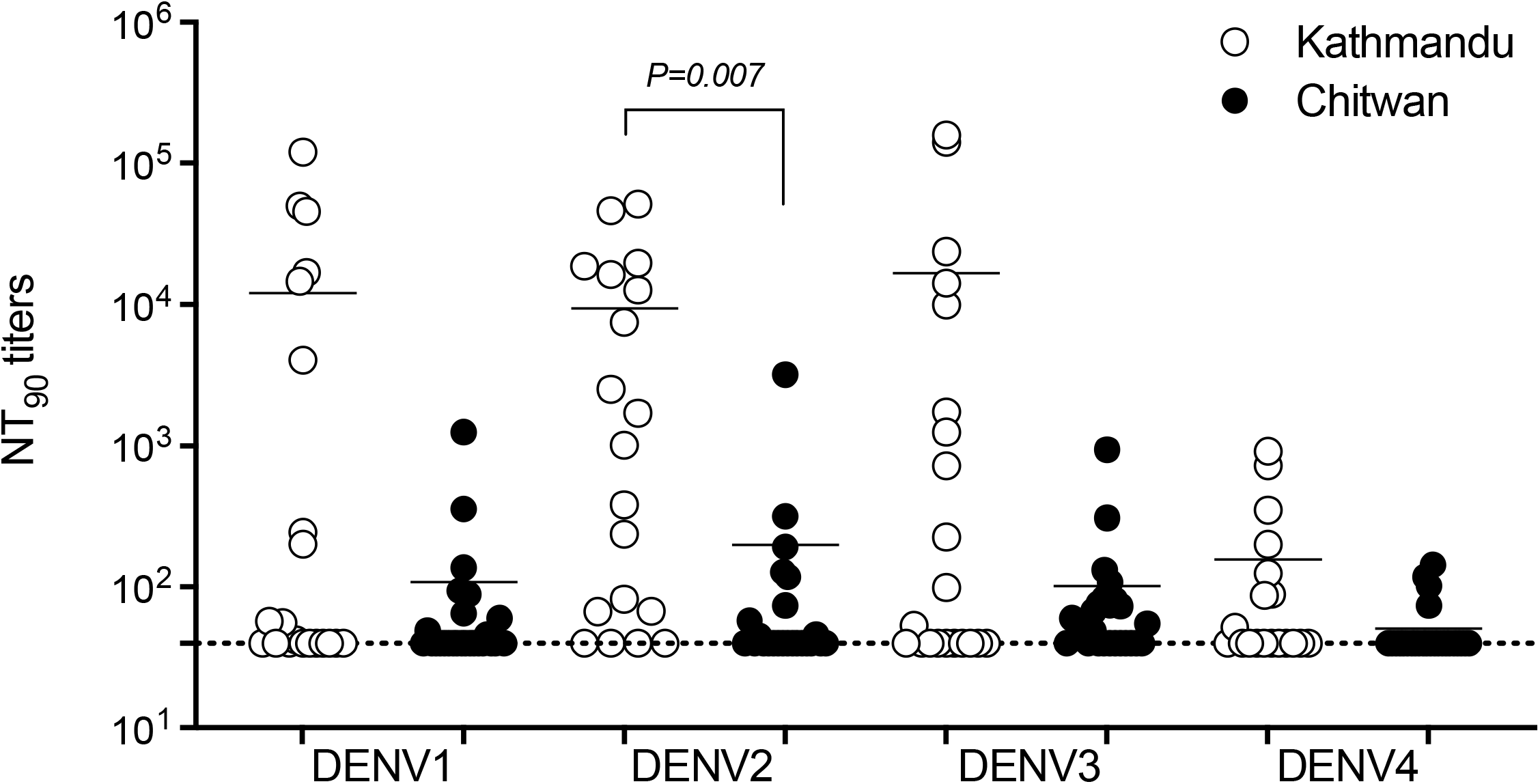
Distribution of nAb titers in samples collected during the 2017 outbreak. **(A and B)** NT_90_ titers were determined according to DENV1–4 reference serotypes (A) and sample collection site (B). Mean values for n=47 (n=21 Kathmandu, n=26 Chitwan) with circles representing individual samples. Dashed lines represent the cutoff value for nAb positivity (NT_90_=40). Mean values were compared using the non-parametric Kruskal–Wallis test.

### Phylogenetic and nucleotide sequence analysis of DENV isolates

Full genome sequences were obtained from the 7 serum samples shown to be positive for DENV RNA by RT-PCR (**Table 2**). Of the 7 genomes, 2 complete sequences and 5 incomplete sequences were obtained. Whole genome phylogenetic analysis of the complete DENV2 genomes (PP152366 and PP152367) indicated that they belong to the Cosmopolitan Genotype, with the closest relatives being DENV2 sequences from Singapore in 2016 and 2014 (MW512465 and MW512428, respectively) (**Fig 3**). Because of the scarcity of full genome sequences from Nepal (n=2), we also analyzed the phylogeny of the E gene of PP152366 and PP152367 andE gene sequences from global DENV2 isolates. Close relationships were detected between PP152366 and PP152367 and E genes from 12 additional DENV2 sequences isolated in 2017 from Nepal (34)) and 4 from India in 2017 (**S1 Fig**). Thus, the 2017 DENV2 strains from Nepal and India likely originated from Singapore.

**Fig 3.**
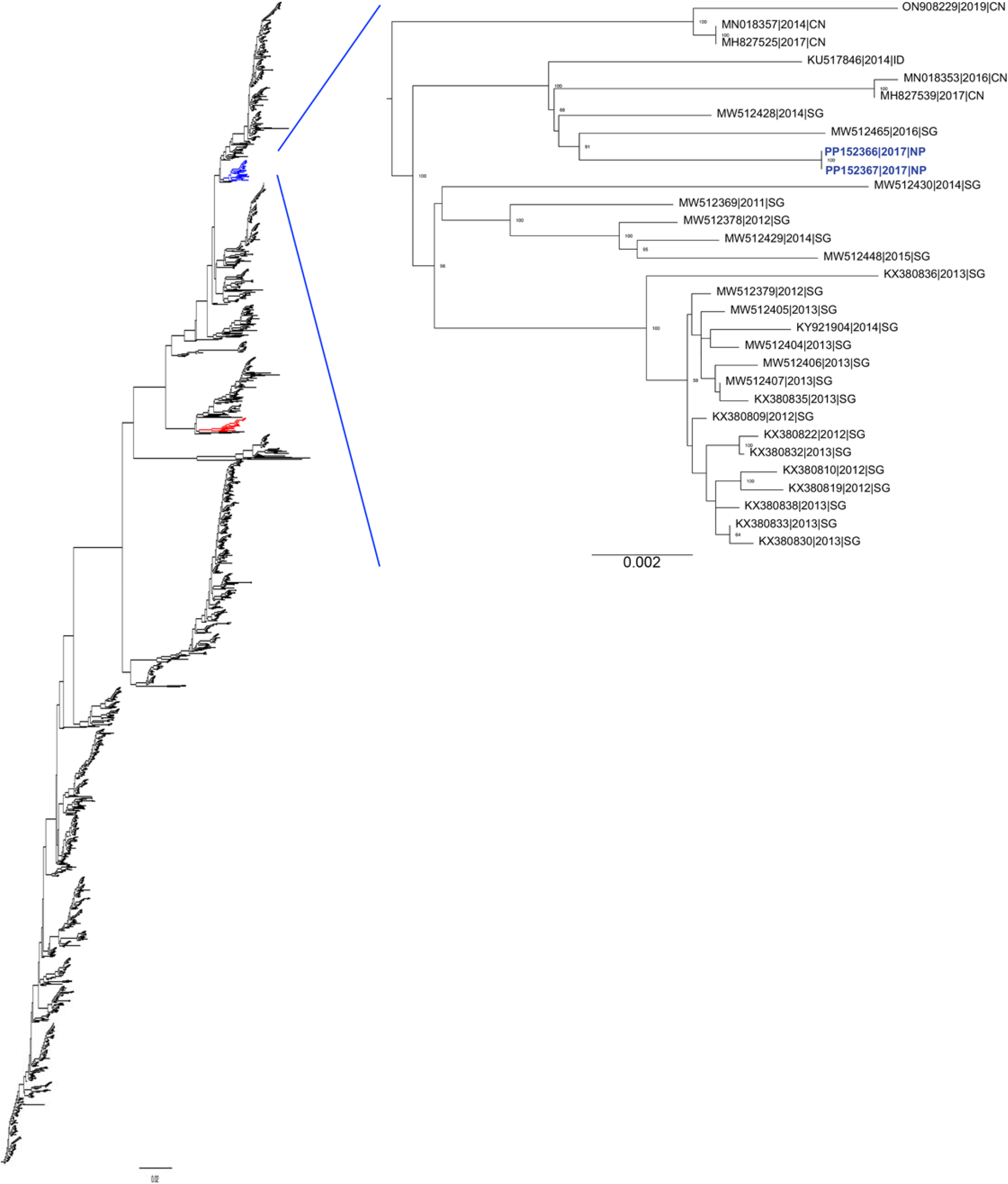
Phylogenetic analysis of full genome sequences of the Nepal 2017 DENV2 isolates generated here and other DENV2 isolates. Phylogenetic tree comparing the DENV2 isolates from this study (blue, PP152366|2017|NP and PP152367|2017|NP), with other DENV2 full genome sequences present in NCBI. Strains are labeled by GenBank ID, as well as year and country of isolation (CN, China; SG, Singapore; NP, Nepal).

## Discussion

Over the past decade, Nepal has experienced increasingly severe outbreaks of dengue in approximately 3-year cycles, and cases were reported from all 77 districts during the 2019 outbreak (25, 35, 36). Despite this public health burden, little was known about the DENV genotypes and serotypes circulating during each outbreak, or of the DENV serological status of the population. This was in large part because the physical infrastructure and knowledge base needed to collect and analyze such data in a systematic manner were lacking, and this study was designed to begin this process with local scientists/clinicians. The overall goal is to amass sufficient data to enable local and central governments to plan for future dengue outbreaks and, eventually, the selection and distribution of DENV vaccines.

We report here the analysis of samples collected during the 2017 dengue season from 49 patients presenting at the main hospitals in Kathmandu and Chitwan in central and southern Nepal, respectively. Most of the cohort were male which is consistent with the other clinical studies of the 2017 dengue season (34, 37) as well as the major DENV outbreaks since 2006 (22, 25, 35, 38, 39). The elevated proportion of men in these cohorts could be because men are more likely than women to engage in healthcare-seeking behaviors and access healthcare, and to have jobs that put them at higher risk for exposure to *Aedes* mosquitos.

This is the first study to examine the DENV-neutralizing capacity in individuals with suspected dengue in Nepal. We selected a higher cutoff for positivity (NT_90_ *vs* standard NT_50_) based on the potential for patients to harbor cross-reactive nAbs (40) arising from previous exposure to other DENV serotypes. Our finding that most of the patient samples did not contain DENV4-specific nAbs is consistent with previous reports demonstrating only a minor contribution of DENV4 to previous DENV outbreaks in Nepal (22, 25, 37). In contrast, DENV1 and DENV2 have been the historically dominant serotypes during major outbreaks, alternating between DENV1 in 2010 and 2016 (41), and DENV2 in 2013 and 2019 (20, 36, 42). In the seven samples serotyped by RT-PCR, six were DENV2 and one was DENV3, suggesting that the dominant circulating DENV serotype may oscillate even in the years between outbreaks. Given that pre-existing Ab responses can limit dengue disease severity (43), and increasing evidence indicates a critical role for serotype-specific nAbs in protecting against infection (44), our data suggest that a DENV4 outbreak in Nepal could lead to more severe outcomes.

Our whole genome sequence analysis was limited by the incomplete sequences obtained from 5 of the 7 DENV RNA+ samples. Phylogenetic analysis of the two complete DENV2 sequences revealed a close relationship with strains from the 2016 dengue outbreak in Singapore (45). Further analysis of E gene sequences showed close relationships with additional DENV strains obtained from Nepal and India in 2017. A previous study also identified between the major DENV isolates circulating in Nepal in 2017 and strains from Singapore (2014, 2016), China (2016, 2017) and Indonesia (2014) (34). Thus, our whole genome and E gene sequence data suggest that the DENV2 2017 Nepal lineage has been circulating in South Asia for at least 5 years. Interestingly, DENV RNA was detected by RT-PCR in only 7 of the 21 samples designated NS1+ by ELISA. This could be explained by NS1 protein having a longer serum half-life compared with viral RNA (46-48), by the presence of DENV RNA mutations that are not complementary to the RT-PCR primers used (49), and/or deterioration of RNA quality during international transportation. The last concern provides an additional justification for our goal to establish the necessary infrastructure in Nepal to surveil and analyze DENV infections within Nepal.

In conclusion, despite its limited size, our study not only sheds light on the prevalence of DENV serotypes and nAbs in a Nepalese cohort during the 2017 dengue season, but also marks a significant advance in establishing, for the first time, the scientific infrastructure to perform genomic and immunologic surveillance of DENV in Nepal. Our analysis demonstrates that a DENV2 genome closely related to strains from Singapore in 2016 was most likely the dominant serotype circulating in Nepal in 2017. The shift in dominant DENV serotypes between outbreaks and the prevalence of both primary and secondary DENV infections underscore the need for sustained surveillance of DENV at virologic and immunologic levels. As noted earlier, the clinical outcome of DENV vaccination will be influenced by the vaccine efficacy profile, the DENV strains circulating at the time of vaccination, and the subjects’ exposure prior to and after vaccination. Thus, our results represent a springboard for further studies to understand DENV virology and immunology across Nepal and thereby inform the selection of one or more DENV vaccine candidates.

### Study limitations

The small cohort size limits the interpretative value of our results. Additionally, complete clinical data were unavailable for some patients. Analysis of samples collected at a single time point during the acute phase of infection, rather than paired acute/convalescent samples, allowed us to infer, but not establish, the subjects’ DENV infection status (50).

## Acknowledgments

We thank the Flow Cytometry Core (Cheryl Kim) at the La Jolla Institute for Immunology for their assistance with experiments, and Dr. Aunji Pradhan from Tribhuvan University for her constructive comments on the manuscript. We are grateful to the clinical staff at Sukraraj Tropical and Infectious Disease Hospital and at Chitwan Medical College and Teaching Hospital, and to the study participants and their families. We would also like to thank Karius Inc. (Redwood City, CA, USA) for its training on regulatory compliance for the source documents and Dr. David Wang (Washington University in St. Louis) NIH grant U01 AI151810) for helping us to build scientific infrastructure at Tribhuvan University.

## Author contributions

SP, AEN, KDM, and SS conceived the study. MMc, AB, SKM, and SRY enrolled patients and collected metadata. SP, JT, MMc, SSh, OMS, and AEN performed the experiments. SP, JT, MMc, AEN, OMS, and RN analyzed and interpreted the data. AEN, MMc, and KDM supervised the experiments. SP, AEN, MMc, OMS, KDM, and SS wrote the manuscript. MY, MLM, KDM, OMS and SS edited the manuscript drafts. All authors approved the final manuscript version.

## Supporting information

**S1 Fig.**
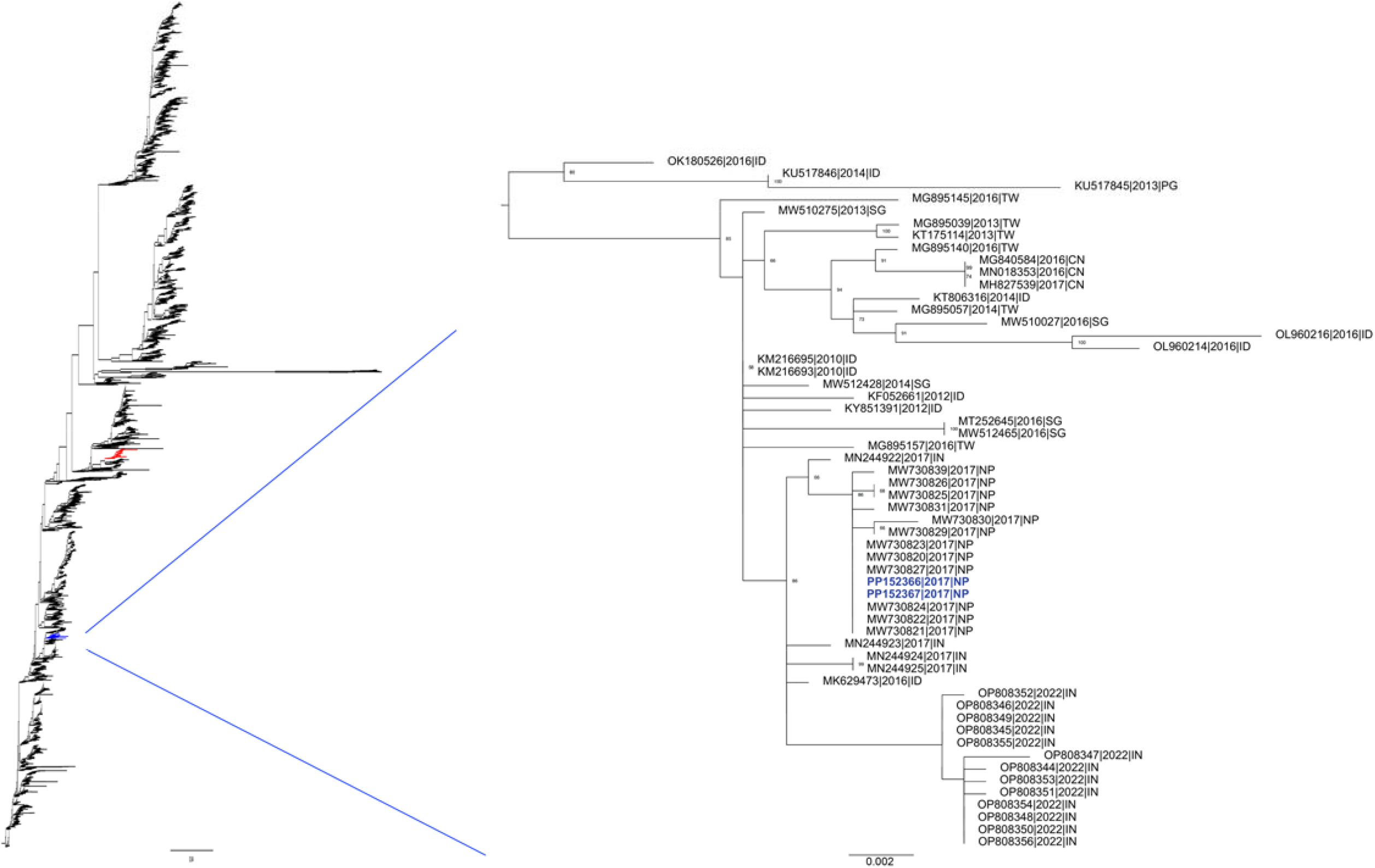
Phylogenetic analysis of E gene sequences from the 2017 Nepalese DENV2 genomes from Nepal obtained in the present study shows a close relationship with other 2017 DENV2 isolates from Nepal and India. Phylogenetic tree comparing E gene sequences of the two 2017 DENV2 strains from Nepal in the current study (blue font, PP152366|2017|NP and PP152367|2017|NP) with other E gene sequences in NCBI. Strains are labeled by GenBank ID, followed by the year and country of isolation (CN, China; ID, Indonesia; IN, India; NP, Nepal; PG, Papua New Guinea; SG, Singapore; TW, Taiwan).

